# The gut microbiota of two rodents varies over fine spatial scales yet is minimally influenced by the environmental microbiota

**DOI:** 10.1101/2025.09.07.673794

**Authors:** ART Figueiredo, KM Wanelik, M Quicray, C Lamberth, A Raulo, SCL Knowles

## Abstract

The gut microbiota of animals varies considerably between individuals within the same population. Given the importance of gut microbiota composition for animal biology and ultimately fitness (e.g., via effects on physiology, immunity and even behaviour), there is considerable interest in understanding which factors shape its variation. Here, we tested whether local environments are an important source of microbial diversity for mammalian guts, by sampling the faecal microbiota of wood mice (*Apodemus sylvaticus*) and bank voles (*Myodes glareolus*), as well as environmental microbiotas, across Wytham Woods. We first find that both species share a low proportion of their microbiotas with local environments and that most shared microbes are aerotolerant. We then detect that microgeographic patterns (namely soil chemistry) are associated with variation in the microbiota of local environments but this does not translate to microbiota variation between animals. Nevertheless, between-individual gut microbiota variation is strongly associated with where animals are sampled, suggesting that other local factors (e.g., biotic environment, interactions between individuals) are much stronger predictors of microbiota composition than local abiotic factors. Finally, we find that wood mice (but not bank voles) are more likely to share microbes with local than non-local environments, but that local environments still contribute little to animal microbiotas. Our findings challenge the prevailing view that mammalian microbiotas are strongly affected by local soil chemistry or the abiotic environment, while further supporting the prediction that biotic and/or other intrinsic factors (which can also act on a local scale) contribute more to microbiota composition.

## Introduction

The mammalian gut microbiota is now known to influence many facets of host biology, from immune development [1] and metabolism [2,3] to pathogen resistance [4] and behaviour [5]. Multiple factors shape the composition of animal gut microbiotas, including within-host factors that affect which of the microbes present in a host are favoured, such as diet [6,7] or host genotype [8,9] and between-host (microbial transmission) processes that affect which microbes colonise host’s epithelial surfaces in the first place [10,11].

Transmission processes are now recognised as a dominant force shaping the gut microbiomes of both wild mammals and humans. Much of this transmission appears to be social in nature, with transmission of gut microbes from other animals and especially conspecifics, being a dominant force shaping the gut microbiota [10,12–14]. But colonising microbes can derive from other sources in a host’s environment, such as soil, food, or other surfaces that hosts come into contact with. Recent studies have suggested this ‘environmental microbiota’ may be an important source of gut colonists, capable of shaping both community composition and host traits. For example, soil microbes have been shown to successfully colonise and persist in the guts of germ-free house mice, influencing the resultant microbiota composition[15,16]. Multiple studies have also cohoused conventional lab mice with soil, and described substantial impacts of soil exposure on gut microbiota composition (comparable in size to dietary effects [17]), with documented knock-on effects for host phenotypes (e.g. immunity[18], recovery of gut microbiota from antibiotics [19]).

However, much of our knowledge about the impact of environmental microbes on the gut microbiota comes from studies in captive animals [20–22] and especially laboratory mice, which are unusual in their domesticated [23,24], inconsistent and easily perturbed gut microbiomes compared to those of wild animals [25,26]. Far less is known about the role of the local environmental microbiota in shaping the gut microbiota of wild mammals in natural habitats. One study in wild baboons found that soil biochemical properties were strong predictors of gut microbiota composition across populations, and much stronger than host-specific factors, but whether this was due to direct colonisation of the gut by soil microbes or indirect effects through diet, was unknown [27]. While several studies have identified fine-scale spatial variation in the gut microbiota of wild mammals[28,29], multiple processes could drive such fine-scale variation besides differences in the environmental microbiota, including spatial variation in diet, or stress, potentially driven by differences in habitat quality [30].

Here, we use a wild rodent study system to investigate the extent to which microbes from the local environment colonise the gut of wild mammals, and whether this explains any spatial variation in gut microbiota composition. We sampled the gut microbiota of two common species of rodent - wood mice (*Apodemus sylvaticus*) and bank voles (*Clethrionomys* [formerly *Myodes*] *glareolus*) as well as the microbiota of environmental substrates in their local environment that they are most likely to come into contact with (soils and leaf litter) from 16 sites across a heterogeneous woodland in which soil chemistry varies widely. As soil chemistry is well known to shape the soil microbiome [31], this study system maximises our chances of detecting fine-scale variation in the soil microbiota and, by extension, of detecting an eventual environment-to-gut transmission signal. By studying mice and voles at the same sites, we compare the strength of environment-to-gut transmission signal across two species with differing habits and ecology; although both species nest in underground burrows dug in soil, nocturnal wood mice are found in open woodland habitat and are more arboreal, while less agile voles are cathemeral and prefer areas of denser understory [32].

## Methods

### Animal trapping

Animal trapping and soil sampling took place at Wytham Woods, Oxford (51°46’N, 1°20’W). To minimise well-documented seasonal variation in the gut microbiota [33], all sampling was conducted in a 2-month period from October to November 2020. To capture as wide a range of environmental microbiomes as possible, 16 sampling plots were selected that covered the breadth of historically classified soil types at Wytham (Fig. S1): (1) limestone, (2) limestone-clay, (3) clay, (4) Corallian sands, (5) clay-sands. Trapping was carried out one night per week in a design where multiple (typically 5) small (50 x 50m) grids were trapped per night with between 20 and 40 traps, in a rotating design with the aim of collecting up to 10 samples per species per grid in total. Small Sherman traps were baited with eight peanuts, a thin slice of apple, and sterile cotton wool for bedding and were set at dusk and collected at dawn the following day. All traps were washed and cleaned with a 1% and then a 20% bleach solution between captures to prevent cross-contamination. To maximise trapping success, traps were pre-baited with peanuts and left clamped open for up to three nights prior to trapping. At first capture, each animal was fur clipped to indicate they had previously been caught, demographic data were recorded (sex, reproductive state, and age class according to body mass and pelage - juvenile, sub-adult, adult). A single faecal sample was collected from the trap for each individual at first capture, kept cool in a fridge during fieldwork and stored at −80^°^C at the end of the day for later gut microbiota analysis. We captured and sampled a total of 134 wood mice (*Apodemus sylvaticus*, AS) and 79 bank voles (*Myodes glareolus*, MG) across 16 sites (Data S1).

### Environmental microbiota sampling

Four environmental microbiota samples were collected per site - leaf litter, surface soil, soil from 5cm and 15cm depth. Environmental samples were collected after the completion of trapping, in December 2020, over a period of 8 days. At each sampling site, samples were collected at 10, approximately equally spaced points, along a ‘W’ shape extending across the 50 x 50m trapping grid. Within a ∼1m area around each sampling point, a leaf litter sample was collected (from a 10 x 10cm area), as well as a surface soil sample (approximately 0.5g), using aseptic techniques. A corer was then pressed into the soil to a depth of 20cm and the resulting core divided into an upper (0-10cm) and a lower (10-20cm) core. Each core was cut in half and a 0.5g sample taken from the middle to acquire a sample at 5cm and 15cm depth for soil microbiota analysis. The remainder of each core was kept for soil chemistry analysis.

For any given site, the 10 collected environmental samples for each of the four depths (litter, surface, 5cm and 15cm) were pooled. For soil microbiota characterisation, sterilised water was added to each pooled soil sample and vortexed to make a homogeneous 1g/ml soil slurry, 300ul of which was added to 300ul DNA/RNA Shield (ZymoBiomics) to achieve a final concentration of 0.5g/ml. Leaf litter samples were pooled in a plastic bag, 200-300ml sterilised water added to each bag before shaking vigorously and finally adding the resulting leaf wash to 300ul of DNA/RNA Shield.

### Soil chemistry analyses

A second set of 10 to 14 soil samples were taken at each site for soil chemistry analyses. Samples of 61cm^3^ (approximately 80g) were collected at depths of 0cm to 10cm and 10cm to 20cm at evenly spaced intervals along the ‘W’ shaped route described above. The core samples from each site were then combined by depth, mixed, stones and twigs removed, and a ∼550g sub-sample retained for analysis. Soil samples were dried at 105ºC and were then ground to <2mm for pH and calcium carbonate concentration determination. We focused on pH and calcium carbonate as soil chemistry variables of interest, as these are known to vary widely across the woodland, with CaCO_3_ variation arising from the fact the woodland sits atop an ancient coral reef [1]. Determination of soil pH in both water and calcium chloride was performed as described in [34,35], using a bench pH meter pH211 (Hanna Instruments Ltd), calibrated at pH 4 and 10 (Hanna Instruments Ltd) using a pH probe (Mettler Toledo™ LE427, Fisher Scientific). Carbonate concentration was performed according to [36], using standard 1M HCl, 1M NaOH and analytical grade CaCl_2_. 2H_2_O and CaCO_3_ (Fisher Scientific) and a bench pH meter were used instead of phenolphthalein indicator because solutions were too cloudy to observe the pH 9 to 9.5 endpoint colour change clearly. To generate a chemical profile of the soil at each site encapsulating all 6 metrics, we performed a PCA analysis with R function *prcomp*. As Principal Component 1 explained 81.9% of variance in soil chemistry (Fig. S2), we retained it for further analysis as the factor describing the chemical properties of soil at each site (henceforth “Soil Chemistry”). PC1 largely reflects an axis relating to pH and CaCO_3_ concentration, with high values indicating a high (alkaline) pH and high CaCO_3_.

### Environment and gut microbiota characterization

The bacterial microbiota of 277 samples was characterized: 134 wood mice and 79 bank voles (each from a unique individual), and 64 environmental (soil or leaf litter) samples. DNA was extracted from faecal and environmental samples using ZymoBiomics DNA Miniprep Kits following the standard kit protocol, and including one pure water extraction control per batch (13 in total). Library preparation and sequencing was performed at the Integrated Microbiome Resource (IMR), Dalhousie University, Canada. The full-length 16S rRNA gene was sequenced using primers 27(F) and 1492(R) [37] on a PacBio platform, alongside all extraction controls and 4 PCR controls. Raw sequencing data was processed through the DADA2 pipeline [38] (v1.22.0) adjusted for PacBio long-read data [39] to infer amplicon sequence variants (ASVs), with the “pooled” argument set to TRUE to detect rare variants across samples. DADA2 was run independently for each sequencing run. Taxonomy was assigned using the SILVA database (version 138.1). After generation of a *phyloseq* object [40], singleton and doubleton ASVs were removed as well as those assigned as *Mitochondria*, and those not assigned to Phylum level. We then used R package *decontam* [41] applying its “prevalence” method and default parameters to further identify and exclude 45 putative contaminant ASVs.

### Statistical Analysis

All statistical analyses were performed in R, version 4.4.2 [42]. First, to visualise how the microbiota of different sample types (rodent guts, leaf litter, and soil) clustered, we performed non-metric multidimensional scaling (NMDS) on the Jaccard Index between samples. Because soil and leaf litter samples clustered together in the NMDS analysis, from then onwards we pooled them one category (Environmental samples), but retained their depth in a separate factor (litter, surface, 5cm and 15cm) when appropriate. We then assessed how many ASVs were detected in each sample type, and created a proportional Venn diagram using package *eulerr* [43]. To explore the aerotolerance phenotypes of ASVs present in each sample type, we assigned aerotolerance (aerotolerant vs anaerobic) to each ASV based on known characteristics of the genus, either from Bergey’s Manual or other literature sources, using the classification made by Hanski et al in [24].

We used Bayesian regression models for statistical analyses implemented in package *brms* [44]. In a first model, we tested whether ASV abundance (as a proxy for origin) is associated with sample type and aerotolerance phenotype (and their interaction) as fixed factors, and ASV identity as a random factor. In subsequent analysis, to model microbiota similarity among samples, we used dyadic models, where the response was a beta diversity metric, and included fixed effects describing dyad level traits, i.e., for factors whether they are the same or a different level, and for continuous variables, the numerical difference between values for each sample in a pair. A multi-membership random intercept term for the samples in each dyad (Sample A + Sample B) was included to account for the non-independence inherent to this type of dyadic data. For all models, we included as a technical covariate the difference in read depth between samples. To allow for a direct comparison between model estimates, all covariates that did not already naturally range between 0 and 1 were scaled to do so.

### Assessing predictors of environmental microbiota similarity

We first tested whether environmental microbiota similarity was predicted by soil, geographical or technical properties. Microbiota similarity among environmental samples (Jaccard Index, the proportion of ASVs in a sample dyad that are shared) was modelled as a function of several soil and geographic predictors: sampling depth (same or different), historically-classified soil type (same or different), difference in soil chemistry PC1, and sampling site (same or different). We also repeated the model replacing sampling site with the Euclidean distance between sampling sites.

### Assessing predictors of gut microbiota similarity

We then tested whether wood mice or bank vole gut microbiota similarity was predicted by intrinsic, geographic, soil, or technical factors. Microbiota similarity (Jaccard Index) was modelled as a function of sex, site and soil type (all as binary factors, same or different), as well as difference in read depth (as previously described). As 6 bank vole dyads had a Jaccard similarity of 0 (no shared ASVs), in this dataset we transformed this metric with a Laplace smoothing so that it would be restricted to range between 0 and 1: Smoothed Jaccard = (*Jaccard* *(*n* − 1) + 0.5) /) [45].

### Assessing predictors of gut-environment microbiota similarity

Finally, we tested whether soil, animal, geographical or technical properties predicted the extent of microbial sharing between wood mice or bank voles and environmental samples. Because a considerable proportion of animal samples did not share any ASVs with environmental samples (Fig. S3), instead of a dyadic model on microbiota similarity, we used a Bernoulli *brms* model with a binary response variable of whether an environment-gut sample pair shared at least one ASV with each other (1), or not (0).

## Results

### Microbial taxa shared between the gut and environmental microbiota

Each sample type (the gut of each rodent species, and environmental samples) contained largely independent sets of microbes (Fig. 1A & B), with shared ASVs typically representing less than 10% or fewer of the ASVs in any given sample type (Fig. 1B). Wood mouse (AS) faecal samples contained on average 240.25 (± 5.57 SE) ASVs, whereas bank vole (MG) samples contained slightly more on average, 275.04 (± 11 SE) ASVs. This contrasted with environmental (Env) samples, in which we found on average 402.75 (± 18.67 SE) ASVs in soil samples, and 1090.38 (± 53.02 SE) ASVs in leaf litter samples. Roughly similar numbers of ASVs were shared between the two rodent gut environments, as were shared between one species of rodent gut and environmental samples (Fig. 1B). Notably, rarefaction curves revealed our sampling to be incomplete especially for environmental samples (Fig. S4). This is a known artifact of running dada2 with the “pool” argument set to *true*, which results in a steep increase in the detection rate of rare ASVs, at the cost of sampling completeness [46]. Both AS and MG samples were dominated by the phyla Firmicutes and Bacteroidota. In environmental samples, Verrucomicrobiota, Proteobacteria and Bacteroidota were the dominant phyla in terms of relative abundance (Fig. S5A). While both rodent’s gut microbiotas were dominated by members of the Lachnospiraceae and Muribaculaceae families, environmental samples were much more taxonomically diverse at the family level (Fig. S5B). ASV diversity (regardless of abundance) in each sample revealed broadly the same patterns (Fig. 1C). ASVs shared between animal and environmental samples mostly belonged to the families Xanthobacteraceae, Chthoniobacteraceae, Muribaculaceae and Lactobacillaceae. The former two are typically found in soils and non-animal environments [47,48], whereas the latter two are commonly associated with rodent guts [49,50].

**Figure 1.**
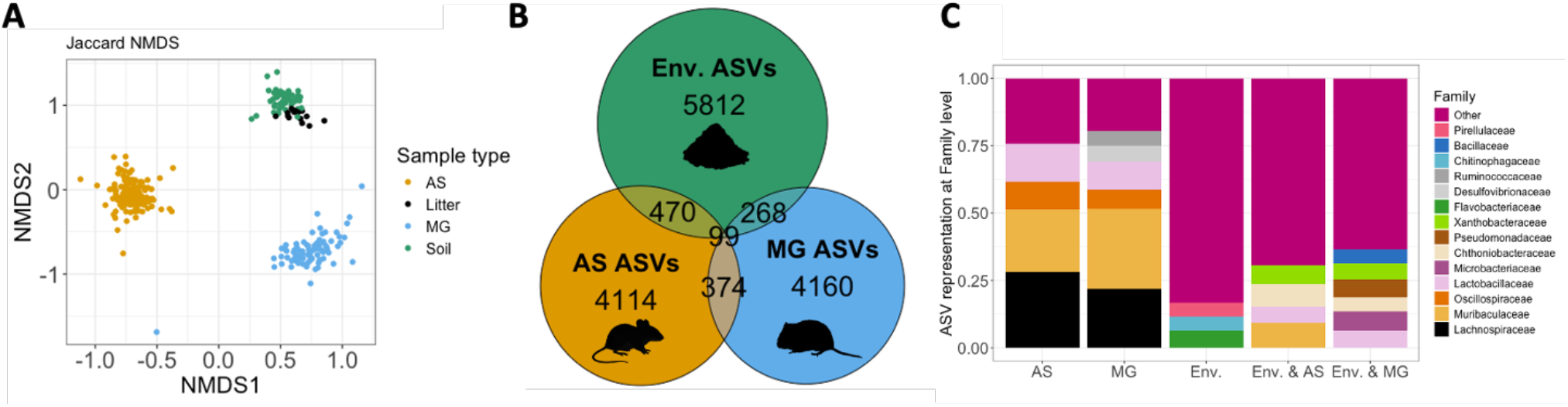
Microbial taxa shared between the gut and environmental microbiota. **A)** Non-metric multidimensional scaling (NMDS) plot of AS, MG, soil and leaf little samples based on Jaccard similarity. **B)** Proportional Venn diagram depicting the total number of distinct ASVs found in each sample type, and the number of ASVs shared between them. **C)** Stacked bar plot depicting the most prevalent microbial families in each sample type (AS, MG and Env), or existing in both gut and environmental samples (Env & AS, Env & MG). Only families containing at least 5% ASVs in each set were named, with the remainder being grouped as “Other”. Stacked bars indicate the proportion of ASVs affiliated with each family in each sample type, or pair of sample types, not relative abundance.

### Taxa shared between the gut and environmental microbiotas are mostly aerotolerant

Of the 544 analysed genera, 181 were classified as aerotolerant, 56 as non-aerotolerant, and 357 as unknown. Further analyses focused only on genera with a known aerotolerance phenotype. Rodent faeces and environmental samples differed in the aerotolerance phenotype of their component taxa. As expected, ASVs detected in rodent faeces largely belonged to anaerobic genera while ASVs detected in environmental samples were overwhelmingly aerotolerant (Fig. 2A). ASVs shared between rodent faeces and the environment mirrored the aerotolerance profile of environmental microbes, being almost entirely aerotolerant (AS & Env = 93% aerotolerant; MG & Env = 96% aerotolerant, Fig. 2A). We found a negative interaction term between sample type and aerotolerance phenotype for ASVs found in the environment and either animal (AS: post. mean = −1.87, 95% CI = −2.41, −1.32; MG: post. mean = −1.91, 95% CI = −3.11, −0.73), suggesting that aerotolerant microbes were significantly more abundant in environmental samples, and non-aerotolerant microbes more abundant when found in animal guts (Fig. 2B & Supp Table 1A).

**Figure 2.**
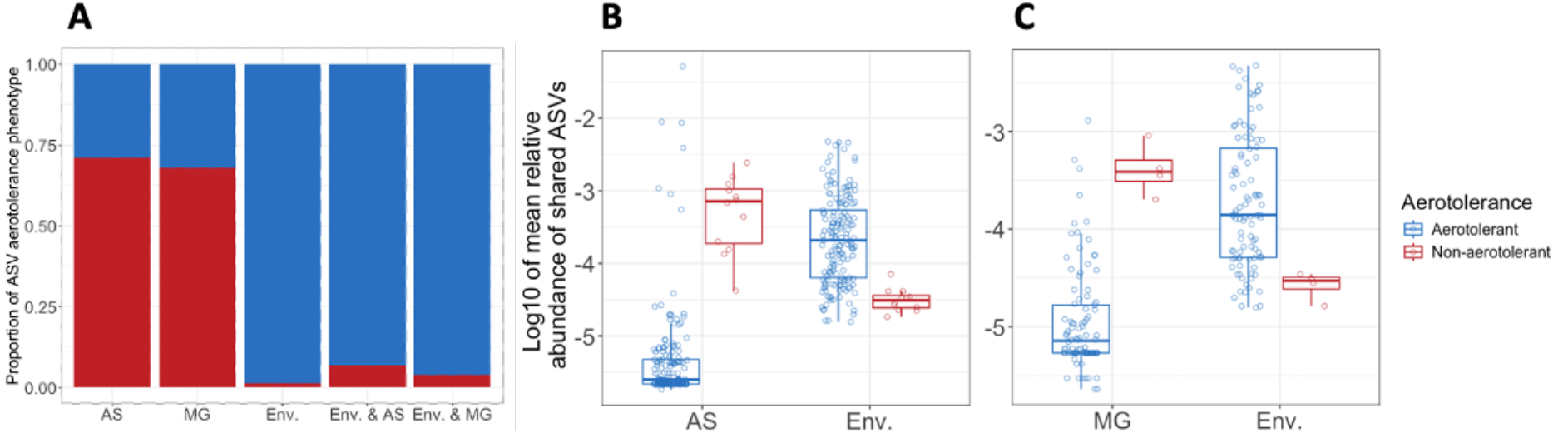
Taxa shared between the gut and environmental microbiotas are mostly aerotolerant. **A)** Stacked bar plot depicting the proportion of ASVs of each aerotolerance phenotype in each sample type, and those found in environmental samples (Env.) *and* either wood mice (AS) or bank voles (MG). Of the ASVs found in both environmental samples and wood mice **(B)** or bank voles **(C)** aerotolerant taxa had higher relative abundance in environmental samples, while non-aerotolerant taxa had higher relative abundance in rodent guts.

### Sampling site and soil chemistry shape the environmental microbiota

Soils from the 16 sites across the woodland varied in several chemical properties, ranging from acidic to moderately alkaline (soil pH range: range 4.40 − 7.85, mean pH = 6.71, sd = 0.92). and varying in soil CaCO_3_ concentration from 0 to 27.7% (mean = 6.33, sd = 7.66). Both geographical and chemical factors predicted variation in environmental microbiota composition (Fig. 3; Supp table 2A). Environmental samples collected at the same sites were more similar to each other than to those collected at different sites (post. mean = 0.25, 95% CI = 0.13, 0.36). Samples of the same type (leaf litter, surface, 5cm or 15cm soil depth) were also more similar than those from different depths (post. mean = 0.31, 95% CI = 0.26, 0.36). Difference in soil chemistry (PC1) across sampling sites also predicted microbiota similarity between samples: the larger the difference in soil chemistry between sites, the fewer ASVs were shared between environmental sample pairs from those sites (post. mean = −0.69, 95% CI = −0.80, −0.56). This result held when sample pairs from the same site were excluded (Fig. S6, Table S2B). The distance between sites did not predict similarity in environmental microbiota among samples better than simply whether or not they were from the same site (“distance” post. mean = −0.09, CI = −0.20, 0.03; Table S3). Historical soil type did not predict microbiota similarity (post. mean = −0.01, 95% CI = − 0.07, 0.05). Read depth difference was accounted for in all models and showed significant associations with microbiota similarity in soils (Fig. S6, Tables S2A and S2B).

**Figure 3.**
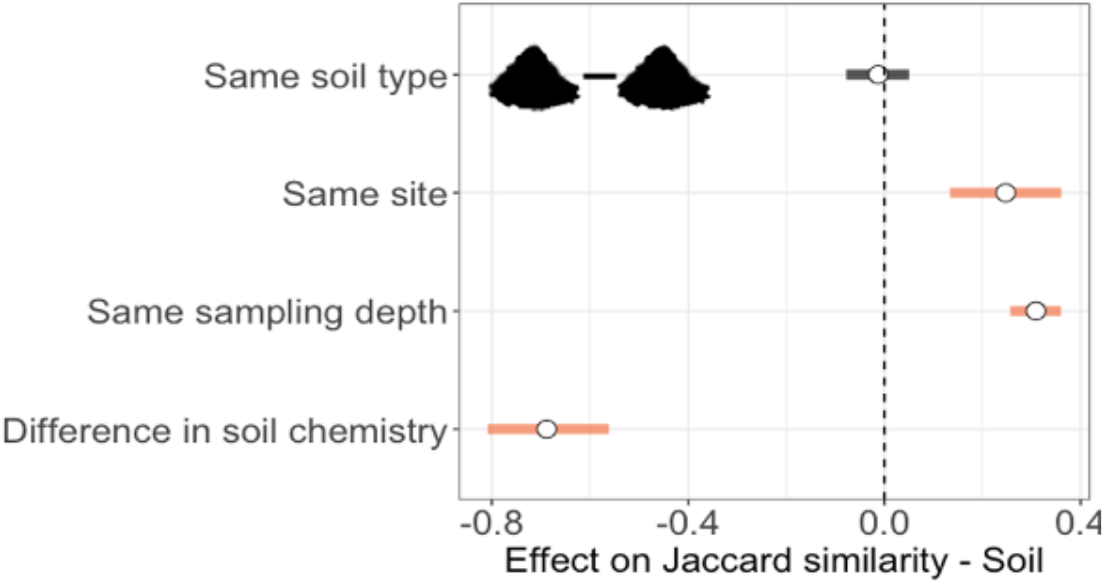
Sampling site and soil chemistry shape the environmental microbiota. The effects of historical soil type, site similarity, sample type/depth similarity, and difference in soil chemistry (y-axis) on environmental microbiota similarity (Jaccard Index, x-axis). Posterior means (white points) and their 95% CIs (lines) obtained from a dyadic Bayesian beta regression model (Table S2A). Orange CI lines do not overlap zero indicating that variable is significantly associated with microbiota similarity, whereas black CI lines overlap zero suggesting that variable does not predict microbiota similarity.

### Gut microbiota of rodents displays fine-scale spatial variation

Both wood mice and bank voles showed similar predictors of gut microbiota similarity (Fig. 4; Table S4). For both species, a clear effect of sampling site was found with conspecifics from the same site sharing more gut microbial taxa than those from different sites (site effect for AS: post. mean = 0.40, 95% CI = 0.34, 0.46; site effect for MG: post. mean = 0.32, 95% CI = 0.22, 0.42). Gut microbiota composition was not predicted by soil chemistry, historical soil type, or host sex (Table S4). Samples taken further apart in time were also less similar in microbiota composition for both species (AS: post. mean = −0.12, 95% CI = −0.19, −0.05; MG: post. mean = −0.40, 95% CI = −0.52, −0.28), consistent with other studies at this site [12,45] and likely due to seasonal shifts in gut microbiota composition [33,51].

**Figure 4.**
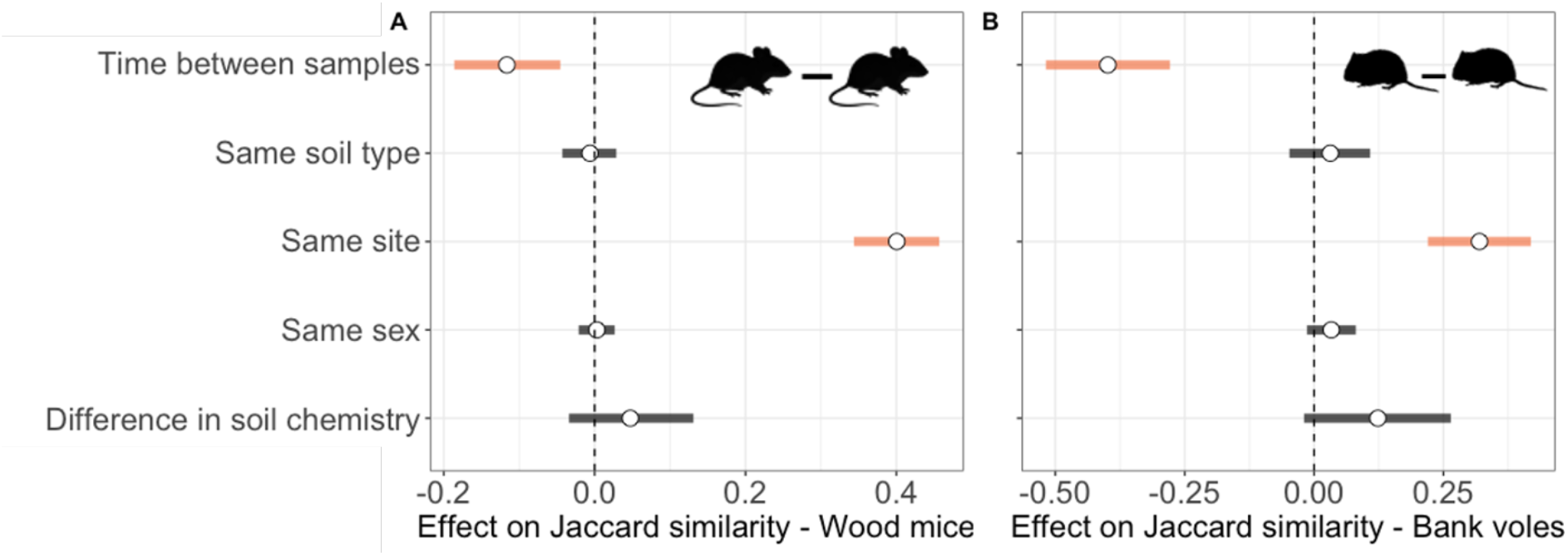
Rodent gut microbiotas show fine-scale spatial variation across the woodland. Predictors (y-axis) of gut microbiota similarity (Jaccard Index, x-axis) among **(A)** wood mice and **(B)** bank voles in Wytham Woods. Posterior means (white points) and their 95% CIs (lines) are plotted from dyadic Bayesian beta regression models (Table S4). Orange CI lines do not overlap zero indicating that variable is significantly associated with microbiota similarity, whereas black CI lines overlap zero suggesting that variable does not significantly predict microbiota similarity.

We found that both wood mouse and bank vole faecal samples were more likely to share ASVs with leaf litter than any other environmental sample type (Fig. 5; AS: post. mean = 1.80, 95% CI = 0.55, 3.13; MG: post. mean = 1.30, 95% CI = 0.17, 2.43; against the reference factor level animal-soil surface; Table S5). In wood mice, we also found that faecal samples were slightly more likely to share ASVs with local than non-local environmental samples (post. mean = 0.43, 95% CI = 0.13, 0.75), and that females were more likely to share ASVs with environmental samples than males (post. mean = 0.84, 95% CI = 0.17, 1.51).

**Figure 5.**
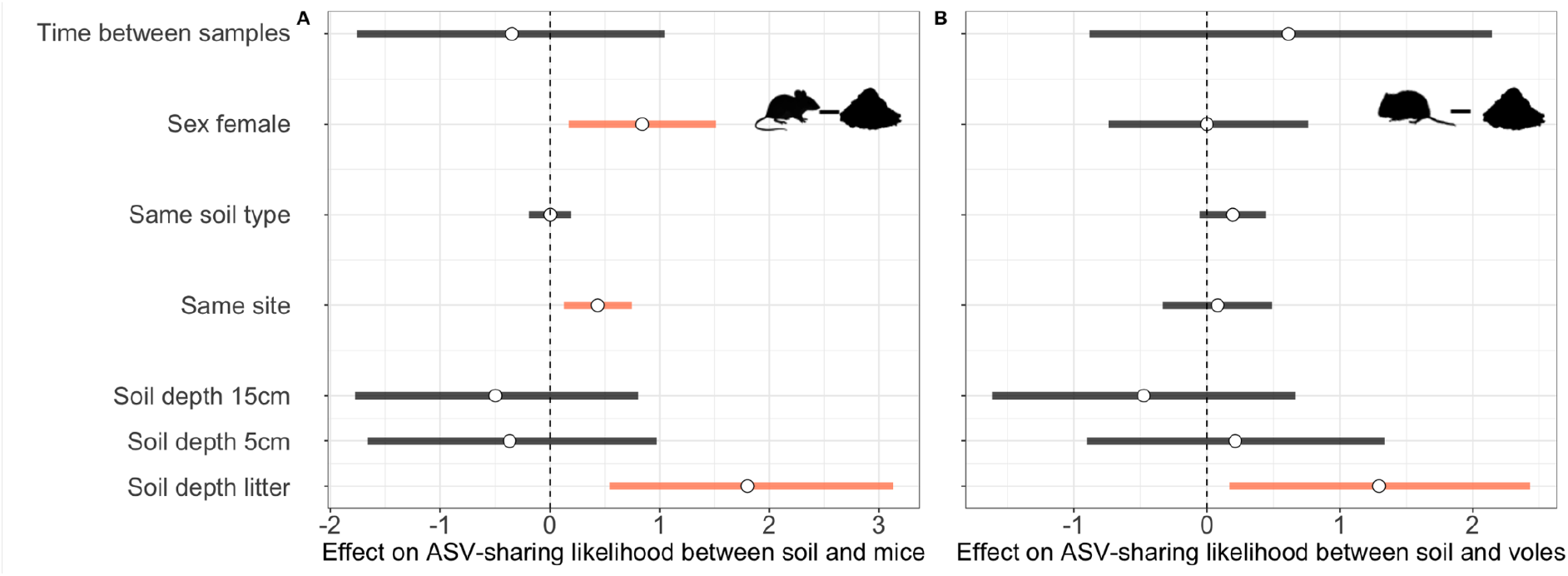
Mouse but not vole gut microbes are more likely to be shared with local than non-local environments. The effects of sampling interval, animal sex, historical soil type, site similarity, and environmental sample type (y-axis) on the likelihood of environmental samples and animal faecal samples (**[A]** wood mice; **[B]** bank voles) sharing at least one ASV (x-axis). Posterior means (white points) and their 95% CIs (lines) are plotted from dyadic Bayesian beta regression models (Table S5). Orange CI lines do not overlap zero, indicating that variable is significantly associated with ASV-sharing likelihood, whereas black CI lines overlap with zero suggesting that variable is not significantly associated with ASV-sharing likelihood.

## Discussion

An important gap in knowledge concerns exactly how important environmental transmission of microbes is for shaping animal gut microbiotas, with studies ranging from arguing it is crucial [17,52,53], to negligible [54–56]. Here, using long-read PacBio sequencing to achieve high taxonomic resolution, we tested the extent to which microbial taxa from relevant environmental substrates are found in the guts of two wild rodent species, and ask whether local environmental microbes might contribute to within-population spatial variation in the rodent gut microbiota.

We first investigated how many gut microbes in wood mice and bank voles were shared with environmental samples. About 10% ASVs found in either animal were also present in environmental samples (Fig. 1B). Most notably, ASVs belonging to Families Lactobacillaceae and Xanthobacteraceae were found in all three sample types (Fig. 1C). The former are known to be associated with animal hosts and contribute to fermentation in the gut [49] while the latter are primarily associated with fresh water and wet soil environments [48]. Wood mice and environmental samples further shared ASVs belonging to the gut-associated [50] family Muribaculaceae and the understudied but soil-associated [47] family Chthoniobacteraceae. Bank voles and the environment shared ASV from the families Bacillaceae, Microbacteriaceae and Pseudomonadaceae. While the former two are ubiquitous and may be environmental- or host-associated [57,58], pseudomonads are mostly found in soils [59]. We furthermore found that the microbes shared between soils and animals are overwhelmingly aerotolerant (Fig. 2A), and more abundant in soils, while the few shared anaerobic microbial taxa are more abundant in animal samples. (Fig. 2B & Fig. 2C) This result is expected, as microbes that can colonize both hosts and the environment will be exposed to oxygen at some point, whereas strictly anaerobic microbes that end up in the environment will likely die soon after. Together, these patterns suggest that microbial transmission between environments and animals is an asymmetrical two-way exchange, with “environmental” microbes occasionally colonising the gut, and persisting at low abundance when they do, while “gut” microbes are found only rarely and at low abundances in environmental substrates. Previous work on wild animals has yielded similar findings: two species of pika (*Ochotona spp*. [56]), tibetan macaques (*Macaca thibetana* [54]), and red-necked stints (*Calidris ruficollis* [55]), share very few ASVs with their surrounding environments.

Despite the small overlap in microbes found in both environmental samples and rodent faecal samples, we still found evidence for fine-scale variation in microbiota composition in both sample types. Environmental microbiota variation was strongly associated with soil chemistry and microgeographic patterns (depth and location) (Fig. 3). Our measure of soil chemistry included both pH and CaCO_3_ concentration, and variation in both these chemical factors has been previously shown to shift soil microbial community composition [60–63]. The causal direction between soil chemistry and microbiota composition is not always clear. However, as a major driver of CaCO_3_ variation in Wytham Woods is the altitudinal variation in geology, with the woodland sitting atop a Jurassic Corallian formation [64], it is safe to assume that soil chemistry begets microbiota composition and not the reverse. We also found that environmental microbiota variation was partly explained by microgeographic patterns, with soil samples from the same site within the woodland, or from the same depth sharing more microbial taxa. These patterns are also expected, as micro-scale plant diversity, land-use and other factors are known to contribute to changes in soil microbiota composition [63].

We then investigated whether a collection of intrinsic and local environment factors were associated with gut microbiota similarity in the two woodland rodent species. First, we found that individuals captured at the same location within the woodland shared more microbes than those captured at different locations, revealing fine-scale spatial variation in gut microbiota composition across this contiguous woodland (Fig. 4). Differences in soil chemistry across capture locations did not predict gut microbiota similarity, suggesting these cannot explain the spatial gut microbiota variation observed. Rather, other effects may explain why animals share more microbes with local conspecifics. Previous studies from the same woodland that used a subset of the sites sampled here, found that social interaction networks but not relatedness explained a significant portion of gut microbiota variation in wood mice [12], alongside seasonal variation suspected to be due to diet [33]. A previous study on *Papio* baboons found soil chemistry to be a strong predictor of microbiota dissimilarity [27], a result we do not replicate in our rodent system. There may be several reasons why our findings differ. This discrepancy could be due to myriad factors, including: (i) the pH range of soils analysed in either study was different (varying by 5 units in [27] compared to 3 in the present study); (ii) the spatial scales of the two studies differing by orders of magnitued, with our sampling locations relatively close together and likely sampling panmictic populations, while Greineisen et al. sample populations separated by hundreds of kilometres; (iii) we focused on calcium carbonate concentration as it is directly relevant for the soil types at our study location, while Grieneisen at al. analysed the effect of exchangeable sodium in the soil. These two factors very likely contribute differently to microbial community compositions in soils, which could also explain differences in results. Potentially, over greater spatial scales and considering sodium rather than calcium, soil chemistry may have a stronger effect on gut microbiota composition in baboons than the rodents studied here, though whether this effect is indirect (e.g., through driving variation in plant diet) or direct (through environmental transmission), remains unknown.

The current paradigm is that local environments can be key sources of microbes for animal gut microbiotas [17,52,53]. From this, follows the expectation that animals should share more microbes with local, than with non-local, environments. Our study design allowed us to effectively test this hypothesis. While we found that wood mice are slightly more likely to share microbes with local than non-local environments, we found no such effect for bank voles (Fig. 5). Several reasons could explain why our results did not support this expectation. First, both animals could move between our sampling locations, thereby being exposed to different set of microbes thus masking a location effect. We find this to be unlikely as both wood mice [65,66] and bank voles [67,68] have home range sizes on the order of magnitude of a hectare (or 100m x 100m) and extending relatively equidistantly in two-dimensional space. This is far less than the average distance between our study plots (1600 m). Second, because rodents burrow in (and are therefore very exposed to) soil, strong mechanisms of host microbiota control may have evolved, thereby minimizing the likelihood of soil microbes colonizing their guts [16]. Third, simple priority or microbial competition effects could explain these patterns [69–71]. For both species, animals were more likely to share microbes with leaf litter than soil. This could be explained by the use of plant matter in nest-lining, frequent movement through leaf litter, as a by-product of plants being part of both species’ diet, or it could be influenced by defecation patterns if defecation is more common in leaf litter than deeper soil. Extending these analyses to other species with different behavioural and life-history patterns will help clarify how local environments shape mammalian gut microbiota. One study did show that the gut microbiota of ring-tailed lemurs (*Lemur catta*) shared more ASVs with their local than non-local environments, but these comparisons included inter-continental and wild vs captive dyads [72].

While the above findings provide valuable insights on how the microbiota of rodents is shaped by their environment, this study has some limitations that warrant consideration. First, we aimed to obtain higher taxonomic resolution through full-length 16S PacBio sequencing, which came at the cost of sequencing depth, thus potentially limiting our ability to detect low-abundance taxa. Shotgun metagenomic sequencing employed in future work could provide even higher taxonomic resolution while also allowing for a functional profiling of the detected microbes. Second, although we focused on two key soil chemistry metrics known to vary across Wytham Woods, other soil chemical traits that were not investigated could influence microbiome variation (such as heavy metal concentrations, imperviousness or salinity). Finally, our faecal sampling protocol involved picking droppings from traps that were left open in the woodland and exposed to the environment. This could have led to environmental microbes being artificially introduced to faecal samples. However, if that were the case, we should have found a strong signal of local environmental microbes on faecal samples, which we did not.

In conclusion, while we found clear fine-scale spatial variation in gut microbiota composition among rodents in a single woodland, we show that environmental microbes in soil and leaf litter are a relatively weak contributor to shaping this variation. The suggestion that environmental microbes may be important players in the development of mammalian immune systems during early life [73,74], is not necessarily inconsistent with the low prevalence of environmental microbes in adult animals as found here. Low-abundance and environmental microbes can have far-reaching effects in animal physiology and microbiota composition [75–77] especially in the development of mammalian immune systems in early life [18,73]. Local environments may thus still be an important source of (possibly predominantly transient) microbes, that while not contributing substantially to adult microbiota composition, still warrant further investigation in terms of their phenotypic impact.

## Data availability

The sequencing data generated for this work will be found in the European Nucleotide Database (https://www.ebi.ac.uk/ena/), under accession no. PRJEB96458. All animal data and code used can be found on GitHub (https://github.com/art-figueiredo/env-apodemus-myodes).

## Acknowledgments

The authors would like to thank JL for field work assistance, and Eveliina Hanski for sharing the aerotolerance classification of microbes. ARTF was supported by a Swiss National Science Foundation Postdoc.Mobility Fellowship (P500PB_206803). KMW was funded by a University of Surrey Future Fellowship while writing up this work. This work was financially supported by the European Research Council under the European Union’s Horizon 2020 research and innovation programme (Grant agreement No. 851550 - to SCLK).

## References

1. Sanidad KZ, Zeng MY. 2020 Neonatal gut microbiome and immunity. Current Opinion in Microbiology 56, 30–37. (doi:10.1016/j.mib.2020.05.011)

2. Gregor R, Probst M, Eyal S, Aksenov A, Sasson G, Horovitz I, Dorrestein PC, Meijler MM, Mizrahi I. 2022 Mammalian gut metabolomes mirror microbiome composition and host phylogeny. The ISME Journal 16, 1262–1274. (doi:10.1038/s41396-021-01152-0)

3. Besser AC, Manlick PJ, Blevins CM, Takacs-Vesbach CD, Newsome SD. 2023 Variation in gut microbial contribution of essential amino acids to host protein metabolism in a wild small mammal community. Ecology Letters 26, 1359–1369. (doi:10.1111/ele.14246)

4. Kamada N, Chen GY, Inohara N, Núñez G. 2013 Control of pathogens and pathobionts by the gut microbiota. Nat Immunol 14, 685–690. (doi:10.1038/ni.2608)

5. Suzuki TA et al. 2025 Selection and transmission of the gut microbiome alone shifts mammalian behavior. (doi:10.1101/2025.01.21.634013)

6. Baniel A et al. 2021 Seasonal shifts in the gut microbiome indicate plastic responses to diet in wild geladas. Microbiome 9, 26. (doi:10.1186/s40168-020-00977-9)

7. David LA et al. 2014 Diet rapidly and reproducibly alters the human gut microbiome. Nature 505, 559–563. (doi:10.1038/nature12820)

8. Benson AK et al. 2010 Individuality in gut microbiota composition is a complex polygenic trait shaped by multiple environmental and host genetic factors. Proc. Natl. Acad. Sci. U.S.A. 107, 18933–18938. (doi:10.1073/pnas.1007028107)

9. Bonder MJ et al. 2016 The effect of host genetics on the gut microbiome. Nat Genet 48, 1407–1412. (doi:10.1038/ng.3663)

10. Sarkar A et al. 2024 Microbial transmission in the social microbiome and host health and disease. Cell 187, 17–43. (doi:10.1016/j.cell.2023.12.014)

11. Heidrich V, Valles-Colomer M, Segata N. 2025 Human microbiome acquisition and transmission. Nat Rev Microbiol (doi:10.1038/s41579-025-01166-x)

12. Raulo A, Allen BE, Troitsky T, Husby A, Firth JA, Coulson T, Knowles SCL. 2021 Social networks strongly predict the gut microbiota of wild mice. The ISME Journal 15, 2601–2613. (doi:10.1038/s41396-021-00949-3)

13. Robinson CD, Bohannan BJ, Britton RA. 2019 Scales of persistence: transmission and the microbiome. Current Opinion in Microbiology 50, 42–49. (doi:10.1016/j.mib.2019.09.009)

14. Brito IL et al. 2019 Transmission of human-associated microbiota along family and social networks. Nat Microbiol 4, 964–971. (doi:10.1038/s41564-019-0409-6)

15. Liu W, Sun Z, Ma C, Zhang J, Ma C, Zhao Y, Wei H, Huang S, Zhang H. 2021 Exposure to soil environments during earlier life stages is distinguishable in the gut microbiome of adult mice. Gut Microbes 13, 1830699. (doi:10.1080/19490976.2020.1830699)

16. Seedorf H et al. 2014 Bacteria from Diverse Habitats Colonize and Compete in the Mouse Gut. Cell 159, 253–266. (doi:10.1016/j.cell.2014.09.008)

17. Zhou D et al. 2018 Soil is a key factor influencing gut microbiota and its effect is comparable to that exerted by diet for mice. F1000Res 7, 1588. (doi:10.12688/f1000research.15297.1)

18. Ottman N et al. 2019 Soil exposure modifies the gut microbiota and supports immune tolerance in a mouse model. Journal of Allergy and Clinical Immunology 143, 1198–1206.e12. (doi:10.1016/j.jaci.2018.06.024)

19. Li N, Zhang H, Bai Z, Jiang H, Yang F, Sun X, Lu Z, Zhou D. 2021 Soil exposure accelerates recovery of the gut microbiota in antibiotic-treated mice. Environ Microbiol Rep 13, 616–625. (doi:10.1111/1758-2229.12959)

20. Rowe SP, Stott MB, Brett B, Juan PAS, Podolyan A, Dhami MK. 2024 Natal soil consumption shifts gut microbiome in captive kiwi (Apteryx rowi). (doi:10.21203/rs.3.rs-5457783/v1)

21. Vo N, Tsai TC, Maxwell C, Carbonero F. 2017 Early exposure to agricultural soil accelerates the maturation of the early-life pig gut microbiota. Anaerobe 45, 31–39. (doi:10.1016/j.anaerobe.2017.02.022)

22. Hyde ER et al. 2016 The Oral and Skin Microbiomes of Captive Komodo Dragons Are Significantly Shared with Their Habitat. mSystems 1. (doi:10.1128/msystems.00046-16)

23. Bowerman KL, Knowles SCL, Bradley JE, Baltrūnaitė L, Lynch MDJ, Jones KM, Hugenholtz P. 2021 Effects of laboratory domestication on the rodent gut microbiome. ISME Communications 1. (doi:10.1038/s43705-021-00053-9)

24. Hanski E et al. 2025 Wild house mice have a more dynamic and aerotolerant gut microbiota than laboratory mice. BMC Microbiol 25, 204. (doi:10.1186/s12866-025-03937-1)

25. Runge S et al. 2025 Laboratory mice engrafted with natural gut microbiota possess a wildling-like phenotype. Nat Commun 16. (doi:10.1038/s41467-025-60554-2)

26. Rosshart SP et al. 2019 Laboratory mice born to wild mice have natural microbiota and model human immune responses. Science 365, eaaw4361. (doi:10.1126/science.aaw4361)

27. Grieneisen LE, Charpentier MJE, Alberts SC, Blekhman R, Bradburd G, Tung J, Archie EA. 2019 Genes, geology and germs: gut microbiota across a primate hybrid zone are explained by site soil properties, not host species. Proc. R. Soc. B. 286, 20190431. (doi:10.1098/rspb.2019.0431)

28. Goertz S et al. 2019 Geographical location influences the composition of the gut microbiota in wild house mice (Mus musculus domesticus) at a fine spatial scale. PLoS ONE 14, e0222501. (doi:10.1371/journal.pone.0222501)

29. Stothart MR, Greuel RJ, Gavriliuc S, Henry A, Wilson AJ, McLoughlin PD, Poissant J. 2021 Bacterial dispersal and drift drive microbiome diversity patterns within a population of feral hindgut fermenters. Molecular Ecology 30, 555–571. (doi:10.1111/mec.15747)

30. Petrullo L, Santangeli A, Wistbacka R, Husby A, Raulo A. 2024 Habitat quality influences gut microbiota via cortisol in Siberian flying squirrels (Pteromys volans). (doi:10.1101/2024.12.11.627940)

31. Islam W, Noman A, Naveed H, Huang Z, Chen HYH. 2020 Role of environmental factors in shaping the soil microbiome. Environ Sci Pollut Res 27, 41225–41247. (doi:10.1007/s11356-020-10471-2)

32. Buesching CD, Newman C, Twell R, Macdonald DW. 2008 Reasons for arboreality in wood mice Apodemus sylvaticus and Bank voles Myodes glareolus. Mammalian Biology 73, 318–324. (doi:10.1016/j.mambio.2007.09.009)

33. Marsh KJ et al. 2022 Synchronous Seasonality in the Gut Microbiota of Wild Mouse Populations. Front. Microbiol. 13, 809735. (doi:10.3389/fmicb.2022.809735)

34. Environment and Heritage - New South Wales. In press. SOIL SURVEY STANDARD TEST METHOD pH: 1:5 SOIL:WATER SUSPENSION.

35. Environment and Heritage - New South Wales. In press. SOIL SURVEY STANDARD TEST METHOD pH: 1:5 SOIL:0.01M CaCL2 SUSPENSION.

36. Food And Agriculture Organization of the United Nations. 2020 Standard operating procedure for soil calcium carbonate equivalent. Titrimetric method.

37. Paliy O, Kenche H, Abernathy F, Michail S. 2009 High-Throughput Quantitative Analysis of the Human Intestinal Microbiota with a Phylogenetic Microarray. Appl Environ Microbiol 75, 3572–3579. (doi:10.1128/AEM.02764-08)

38. Callahan BJ, McMurdie PJ, Rosen MJ, Han AW, Johnson AJA, Holmes SP. 2016 DADA2: High-resolution sample inference from Illumina amplicon data. Nat Methods 13, 581–583. (doi:10.1038/nmeth.3869)

39. Callahan BJ, Wong J, Heiner C, Oh S, Theriot CM, Gulati AS, McGill SK, Dougherty MK. 2019 High-throughput amplicon sequencing of the full-length 16S rRNA gene with single-nucleotide resolution. Nucleic Acids Research 47, e103–e103. (doi:10.1093/nar/gkz569)

40. McMurdie PJ, Holmes S. 2013 phyloseq: An R Package for Reproducible Interactive Analysis and Graphics of Microbiome Census Data. PLoS ONE 8, e61217. (doi:10.1371/journal.pone.0061217)

41. Davis NM, Proctor DM, Holmes SP, Relman DA, Callahan BJ. 2018 Simple statistical identification and removal of contaminant sequences in marker-gene and metagenomics data. Microbiome 6, 226. (doi:10.1186/s40168-018-0605-2)

42. R Core Team. 2024 R: A Language and Environment for Statistical Computing R Foundation for Statistical Computing.

43. Larsson J, Godfrey AJR, Gustafsson P, algorithms) DHE (geometric, code) EH (root solver, Privé F. 2024 eulerr: Area-Proportional Euler and Venn Diagrams with Ellipses.

44. Bürkner P-C. 2017 brms: An R Package for Bayesian Multilevel Models Using Stan. J. Stat. Soft. 80. (doi:10.18637/jss.v080.i01)

45. Raulo A et al. 2024 Social and environmental transmission spread different sets of gut microbes in wild mice. Nat Ecol Evol 8, 972–985. (doi:10.1038/s41559-024-02381-0)

46. Kleine Bardenhorst S, Vital M, Karch A, Rübsamen N. 2022 Richness estimation in microbiome data obtained from denoising pipelines. Computational and Structural Biotechnology Journal 20, 508–520. (doi:10.1016/j.csbj.2021.12.036)

47. 2015 Chthoniobacteraceae fam. nov. In Bergey’s Manual of Systematics of Archaea and Bacteria, pp. 1–1. Wiley. (doi:10.1002/9781118960608.fbm00256)

48. In press. Xanthobacter. In Bergey’s Manual® of Systematic Bacteriology, pp. 555–566. New York: Springer-Verlag. (doi:10.1007/0-387-29298-5_135)

49. Walter J, O’Toole PW. 2023 Microbe Profile: The Lactobacillaceae. Microbiology 169. (doi:10.1099/mic.0.001414)

50. Lagkouvardos I et al. 2019 Sequence and cultivation study of Muribaculaceae reveals novel species, host preference, and functional potential of this yet undescribed family. Microbiome 7. (doi:10.1186/s40168-019-0637-2)

51. Maurice CF, Knowles SCL, Ladau J, Pollard KS, Fenton A, Pedersen AB, Turnbaugh PJ. 2015 Marked seasonal variation in the wild mouse gut microbiota. The ISME Journal 9, 2423–2434. (doi:10.1038/ismej.2015.53)

52. Roslund MI, Laitinen OH, Sinkkonen A. 2024 Scoping review on soil microbiome and gut health— Are soil microorganisms missing from the planetary health plate? People and Nature 6, 1078–1095. (doi:10.1002/pan3.10638)

53. Schnorr SL. 2020 The soil in our microbial DNA informs about environmental interfaces across host and subsistence modalities. Philosophical Transactions B

54. Xu X, Xia Y, Sun B. 2022 Linking the bacterial microbiome between gut and habitat soil of Tibetan macaque (Macaca thibetana). Ecology and Evolution 12. (doi:10.1002/ece3.9227)

55. Risely A, Waite D, Ujvari B, Klaassen M, Hoye B. 2017 Gut microbiota of a long-distance migrant demonstrates resistance against environmental microbe incursions. Molecular Ecology 26, 5842–5854. (doi:10.1111/mec.14326)

56. Li H, Li T, Yao M, Li J, Zhang S, Wirth S, Cao W, Lin Q, Li X. 2016 Pika Gut May Select for Rare but Diverse Environmental Bacteria. Front. Microbiol. 7. (doi:10.3389/fmicb.2016.01269)

57. Mandic-Mulec I, Stefanic P, Van Elsas JD. 2015 Ecology of Bacillaceae. Microbiol Spectr 3, 3.2.16. (doi:10.1128/microbiolspec.TBS-0017-2013)

58. Evtushenko LI. 2015 Microbacteriaceae. In Bergey’s Manual of Systematics of Archaea and Bacteria (ed WB Whitman), pp. 1–14. Wiley. (doi:10.1002/9781118960608.fbm00035)

59. Mehnaz S, editor. 2017 Rhizotrophs: Plant Growth Promotion to Bioremediation. Singapore: Springer Singapore. (doi:10.1007/978-981-10-4862-3)

60. Coller E, Cestaro A, Zanzotti R, Bertoldi D, Pindo M, Larger S, Albanese D, Mescalchin E, Donati C. 2019 Microbiome of vineyard soils is shaped by geography and management. Microbiome 7, 140. (doi:10.1186/s40168-019-0758-7)

61. Rousk J, Bååth E, Brookes PC, Lauber CL, Lozupone C, Caporaso JG, Knight R, Fierer N. 2010 Soil bacterial and fungal communities across a pH gradient in an arable soil. The ISME Journal 4, 1340–1351. (doi:10.1038/ismej.2010.58)

62. Guo A, Ding L, Tang Z, Zhao Z, Duan G. 2019 Microbial response to CaCO3 application in an acid soil in southern China. Journal of Environmental Sciences 79, 321–329. (doi:10.1016/j.jes.2018.12.007)

63. Fierer N. 2017 Embracing the unknown: disentangling the complexities of the soil microbiome. Nat Rev Microbiol 15, 579–590. (doi:10.1038/nrmicro.2017.87)

64. Savill PS, editor. 2011 The Physical Environment. In Wytham Woods: Oxford’s ecological laboratory, Oxford: Oxford University Press. (doi:10.1093/acprof:osobl/9780199605187.001.0001)

65. Benhamou S. 1991 An analysis of movements of the wood mouse Apodemus sylvaticus in its home range. Behavioural Processes 22, 235–250. (doi:10.1016/0376-6357(91)90097-J)

66. Attuquayefio DK, Gorman ML, Wolton RJ. 1986 Home range sizes in the Wood mouse Apodemus sylvaticus: habitat, sex and seasonal differences. Journal of Zoology 210, 45–53. (doi:10.1111/j.1469-7998.1986.tb03619.x)

67. Koskela E, Mappes T, Ylonen H. 1997 Territorial Behaviour and Reproductive Success of Bank Vole Clethrionomys glareolus Females. The Journal of Animal Ecology 66, 341. (doi:10.2307/5980)

68. Kozakiewicz M, Chołuj A, Kozakiewicz A. 2007 Long-distance movements of individuals in a free-living bank vole population: an important element of male breeding strategy. Acta Theriol 52, 339–348. (doi:10.1007/BF03194231)

69. Sprockett D, Fukami T, Relman DA. 2018 Role of priority effects in the early-life assembly of the gut microbiota. Nat Rev Gastroenterol Hepatol 15, 197–205. (doi:10.1038/nrgastro.2017.173)

70. Debray R, Herbert RA, Jaffe AL, Crits-Christoph A, Power ME, Koskella B. 2022 Priority effects in microbiome assembly. Nat Rev Microbiol 20, 109–121. (doi:10.1038/s41579-021-00604-w)

71. Figueiredo ART, Kramer J. 2020 Cooperation and Conflict Within the Microbiota and Their Effects On Animal Hosts. Front. Ecol. Evol. 8, 132. (doi:10.3389/fevo.2020.00132)

72. Bornbusch SL, Greene LK, Rahobilalaina S, Calkins S, Rothman RS, Clarke TA, LaFleur M, Drea CM. 2022 Gut microbiota of ring-tailed lemurs (Lemur catta) vary across natural and captive populations and correlate with environmental microbiota. anim microbiome 4. (doi:10.1186/s42523-022-00176-x)

73. Mulder IE et al. 2009 Environmentally-acquired bacteria influence microbial diversity and natural innate immune responses at gut surfaces. BMC Biol 7, 79. (doi:10.1186/1741-7007-7-79)

74. Donald K, Finlay BB. 2023 Early-life interactions between the microbiota and immune system: impact on immune system development and atopic disease. Nat Rev Immunol 23, 735–748. (doi:10.1038/s41577-023-00874-w)

75. Han G, Vaishnava S. 2023 Microbial underdogs: exploring the significance of low-abundance commensals in host-microbe interactions. Exp Mol Med 55, 2498–2507. (doi:10.1038/s12276-023-01120-y)

76. Han G, Luong H, Vaishnava S. 2022 Low abundance members of the gut microbiome exhibit high immunogenicity. Gut Microbes 14, 2104086. (doi:10.1080/19490976.2022.2104086)

77. Benjamino J, Lincoln S, Srivastava R, Graf J. 2018 Low-abundant bacteria drive compositional changes in the gut microbiota after dietary alteration. Microbiome 6, 86. (doi:10.1186/s40168-018-0469-5)

