## Supplementary figures and tables for "The gut microbiota of two rodents varies over fine spatial scales yet is minimally influenced by the environmental microbiota"

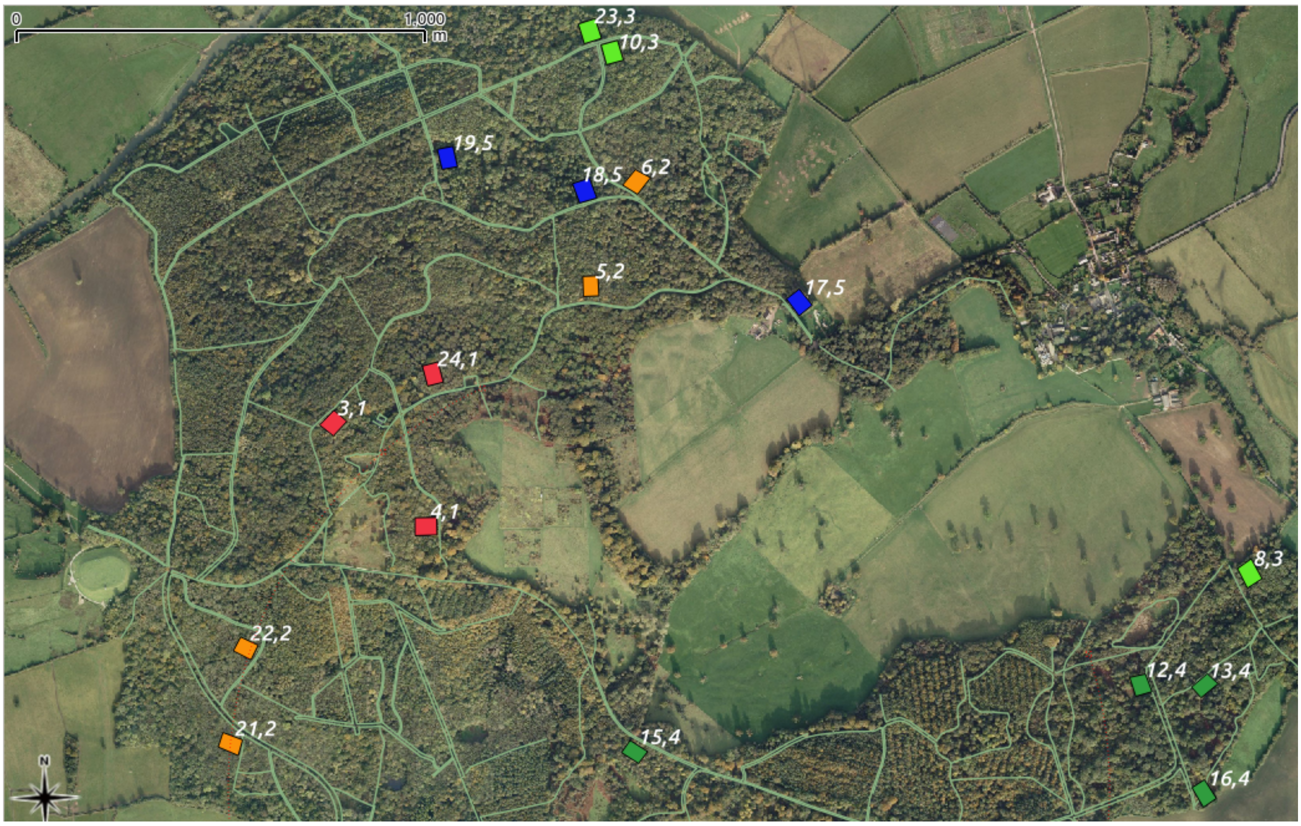


**Fig. S1**: Satellite image of Wytham Woods displaying all study sites where animals were trapped and environmental samples collected. The first number of each site depicts its individual site ID, whereas the second number and color depict its historical soil type classification: (1;red) limestone, (2;orange) limestone-clay, (3; light green) clay, (4;dark green) Corallian sands, (5;blue) clay-sands. Site "12,4" was excluded from further analysis as no animals were captured there.

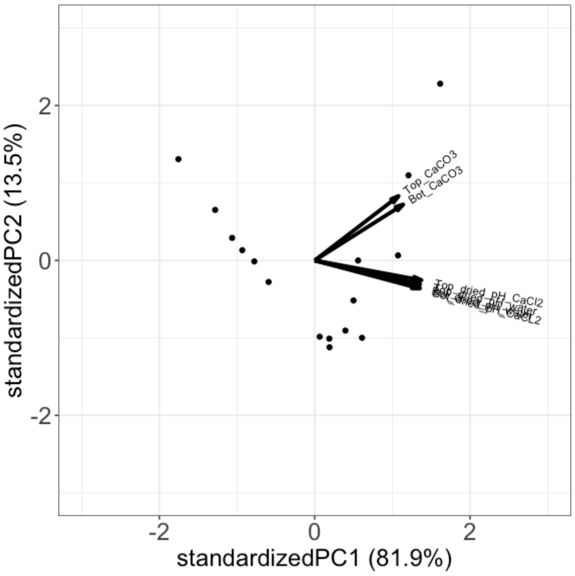


**Fig. S2**: Profiling of each soil sample by PCA analysis of the 6 soil chemistry variables: $CaCO_3$ , pH measured in $H_2O$ and in $CaCl_2$ each in top (0-10cm) and button (10-20cm) soil. Principal Component 1 explains 81.9% variation in soil chemistry.

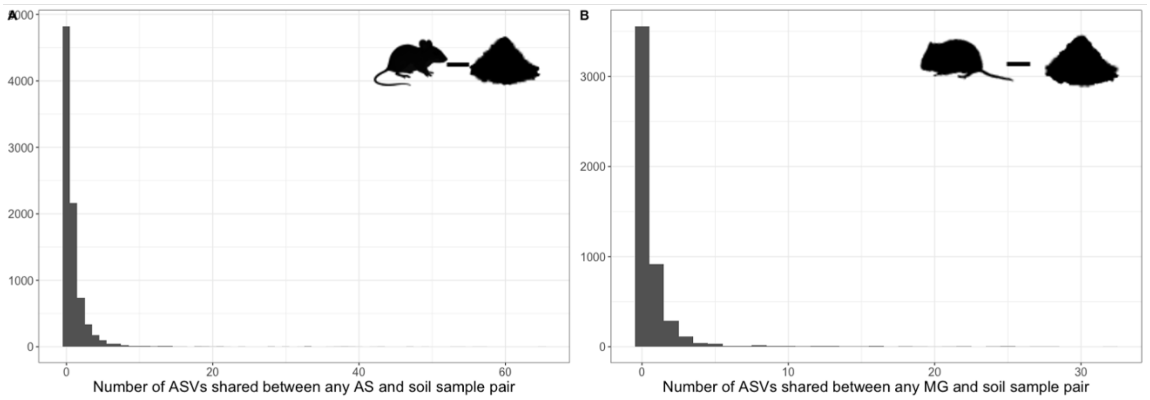


**Fig. S3:** Histogram of frequencies of shared ASVs between any wood mouse - environmental sample pair (A) and bank vole - environmental sample pair (B). Notably, the vast majority of animal-environmental sample dyads shared 0 ASVs.

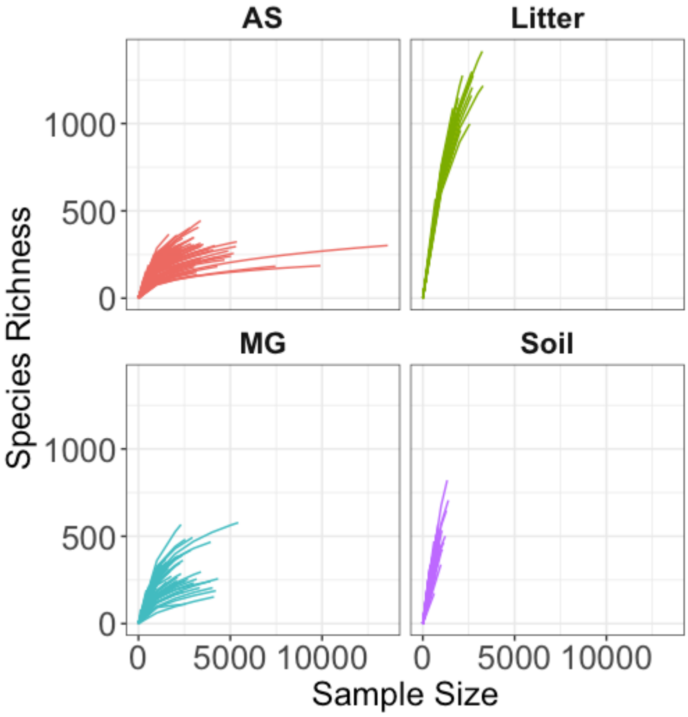


**Fig. S4:** Rarefaction curves for each sample type, before collapsing "soil" and "litter" samples into "environmental" samples. The x-axis depicts sample read depth, whereas the y-axis represents the number of ASVs detected. The low completeness of both litter and soil samples can be explained by the fact that we opted to run dada2 with the "pooled" parameter set to TRUE, which results in many more rare ASVs being retained.

A

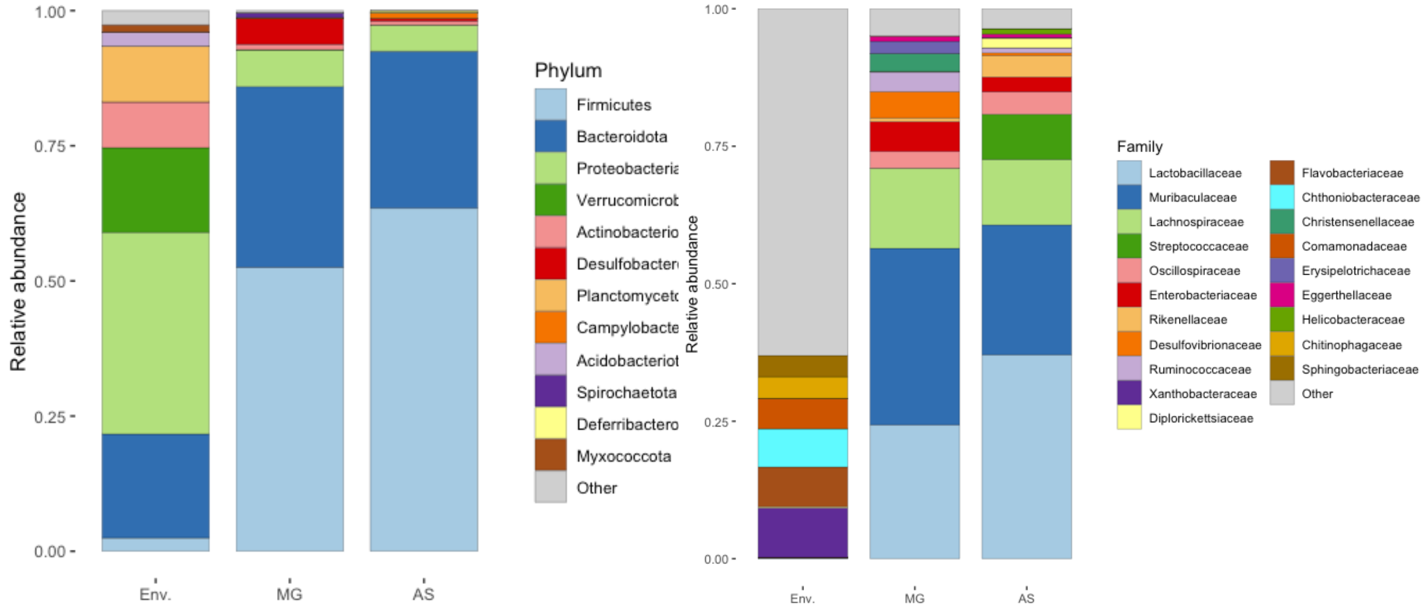


**Fig. S5**: Stacked bar plots depicting the relative abundance of the top 20 microbial phyla (A) and families (B) across each sample type.

A

| Predictor | Estimate | l-95% CI | u-95% CI |
|---|---|---|---|
| Sample type Env. | 0.7999 | 0.6179 | 0.9796 |
| Non-aerotolerant | 1.1761 | 0.7928 | 1.5263 |
| Sample type Env.:Non-aerotolerant | −1.8698 | −2.4100 | −1.3261 |

B

| Predictor | Estimate | l-95% CI | u-95% CI |
|---|---|---|---|
| Sample type Env. | 0.7806 | 0.5828 | 0.9811 |
| Non-aerotolerant | 1.0831 | 0.2101 | 1.7626 |
| Sample type Env.:Non-aerotolerant | −1.9133 | −3.1111 | −0.7388 |

**Table S1:** Results of Bayesian regression (brms) models testing the effect of sample type (Environmental or animal), aerotolerance category (aerotolerant or non-aerotolerant) and their interaction on the relative abundance of shared ASVs. Table A outputs the results for ASVs shared between wood mice and environmental samples, whereas Table B outputs the results for ASVs shared between bank voles and environmental samples. All terms were found to be significant as no 95% credible intervals overlap with zero.

A

| Predictor | Estimate | lCI | uCI |
|---|---|---|---|
| Difference in read depth | −2.3264 | −2.4853 | −2.1698 |
| Difference in soil chemistry | −0.6881 | −0.8089 | −0.5615 |
| Same sampling depth | 0.3089 | 0.2563 | 0.3598 |
| Same site | 0.2471 | 0.1334 | 0.3604 |
| Same soil type | −0.0127 | −0.0777 | 0.0506 |

B

| Predictor | Estimate | lCI | uCI |
|---|---|---|---|
| Difference in read depth | −2.2667 | −2.4315 | −2.1049 |
| Difference in soil chemistry | −0.6416 | −0.7696 | −0.5105 |
| Same sampling depth | 0.3217 | 0.2697 | 0.3745 |
| Same soil type | −0.0107 | −0.0746 | 0.0514 |

**Table S2**: Results of Bayesian regression (brms) models testing predictors of environmental microbiota similarity (Jaccard Index) among environmental sample pairs, either (A) including or (B) excluding sample pairs from the same site.

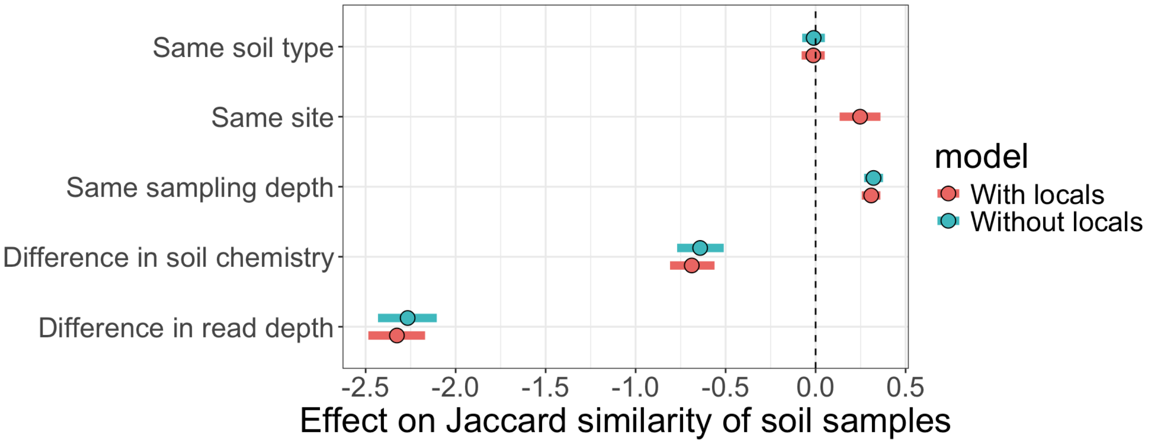


**Fig. S6**: Results of Bayesian regression (brms) models testing the effect of soil type similarity, site similarity (in model with locals), sampling depth similarity, difference in soil chemistry, and difference in read depth on Jaccard similarity between environmental samples. The CIs depicted in red come from a model including all possible environmental sample pairs, whereas the CIs depicted in blue come from a model excluding pairs of environmental samples from the same site. The model outputs are qualitatively similar, suggesting that any eventual co-linearity introduced by sample pairs arising from the same site (as all pairs from the same site will always have the same soil chemistry value) does not affect our results.

| Predictor | Estimate | lCI | uCI |
|---|---|---|---|
| Difference in read depth | −2.3159 | −2.4730 | −2.1575 |
| Difference in soil chemistry | −0.7589 | −0.8789 | −0.6394 |
| Same sampling depth | 0.2972 | 0.2463 | 0.3472 |
| Distance | −0.0887 | −0.2009 | 0.0268 |
| Same soil type | 0.0284 | −0.0325 | 0.0887 |

**Table S3**: Results of brms models testing the effect of predictors on environmental microbiota similarity (Jaccard Index). This model is identical to that for which results are presented in Table S2A, except the binary predictor "same site" is replaced by a continuous predictor "distance".

A

| Predictor | Estimate | lCI | uCI |
|---|---|---|---|
| Difference in read depth | −2.1922 | −2.3759 | −2.0085 |
| Same site | 0.4004 | 0.3436 | 0.4568 |
| Time between samples | −0.1162 | −0.1860 | −0.0454 |
| Difference in soil chemistry | 0.0473 | −0.0340 | 0.1309 |
| Same sex | 0.0029 | −0.0211 | 0.0267 |
| Same soil type | −0.0062 | −0.0428 | 0.0287 |

B

| Predictor | Estimate | lCI | uCI |
|---|---|---|---|
| Difference in read depth | −1.0602 | −1.2412 | −0.8759 |
| Same site | 0.3196 | 0.2199 | 0.4190 |
| Time between samples | −0.3985 | −0.5181 | −0.2784 |
| Difference in soil chemistry | 0.1231 | −0.0195 | 0.2641 |
| Same sex | 0.0333 | −0.0137 | 0.0807 |
| Same soil type | 0.0314 | −0.0477 | 0.1083 |

**Table S4**: Results of Bayesian regression (brms) models testing the effect of difference in read depth, difference in sampling dates, site similarity, difference in soil chemistry, sex similarity, and soil type similarity on wood mice (A), and bank vole B) microbiota similarity (Jaccard Index).

A

| Predictor | Estimate | lCI | uCI |
|---|---|---|---|
| Read Depth difference | −1.9341 | −3.4524 | −0.4099 |
| Time between samples | −0.3461 | −1.7568 | 1.0487 |
| Sex female | 0.8418 | 0.1685 | 1.5104 |
| Same soil type | 0.0038 | −0.1922 | 0.1950 |
| Same site | 0.4348 | 0.1290 | 0.7450 |
| Soil depth 15cm | −0.4973 | −1.7786 | 0.8052 |
| Soil depth 5cm | −0.3672 | −1.6608 | 0.9729 |
| Soil depth litter | 1.7998 | 0.5463 | 3.1256 |

B

| Predictor | Estimate | lCI | uCI |
|---|---|---|---|
| Read Depth difference | −0.4633 | −1.2344 | 0.3046 |
| Time between samples | 0.6128 | −0.8854 | 2.1462 |
| Sex female | −0.0012 | −0.7422 | 0.7634 |
| Same soil type | 0.1935 | −0.0559 | 0.4417 |
| Same site | 0.0813 | −0.3345 | 0.4906 |
| Soil depth 15cm | −0.4749 | −1.6152 | 0.6632 |
| Soil depth 5cm | 0.2123 | −0.9035 | 1.3376 |
| Soil depth litter | 1.2955 | 0.1678 | 2.4341 |

**Table S5:** Results of Bayesian regression (brms) models testing the effect of difference in read depth, time between sampling, sex of animals, historical soil type, site similarity, and the different depths of environmental sampling on the likelihood of wood mice (Table A), and bank vole (Table B) sharing any ASVs with environmental samples.
